# ALS patient-derived motor neuron networks exhibit microscale dysfunction and mesoscale compensation rendering them highly vulnerable to perturbation

**DOI:** 10.1101/2024.01.04.574167

**Authors:** Vegard Fiskum, Nicolai Winter-Hjelm, Nicholas Christiansen, Axel Sandvig, Ioanna Sandvig

## Abstract

Amyotrophic lateral sclerosis affects upper and lower motor neurons, causing progressive neuropathology leading to structural and functional alterations of affected neural networks long prior to development of symptoms. Certain genetic mutations, such as expansions in *C9orf72*, predispose motor neuron populations to pathological dysfunction. However, it is not known how underlying pathological predisposition affects structural and functional dynamics within vulnerable networks. Here, we studied micro-and mesoscale dynamics of ALS patient derived motor neuron networks over time. We show, for the first time, that ALS patient derived motor neurons with endogenous genetic predisposition develop classical ALS cytopathology in the form of cytoplasmic TDP-43 inclusions and self-organise into computationally efficient networks, albeit with functional hallmarks of higher metabolic cost compared to healthy controls. These hallmarks included microscale impairments and mesoscale compensation including increased centralisation of function. Moreover, we show that these networks are highly susceptible to transient perturbation by exhibiting induced hyperactivity.

## Introduction

Amyotrophic lateral sclerosis (ALS) is a neurodegenerative disorder which selectively affects upper and lower motor neurons (MNs), leading to progressive muscle weakness and eventually death by respiratory failure (Masrori and Van Damme 2020, Srinivasan and Rajasekaran 2020). Multiple genetic and environmental factors have been identified, which increase the risk of developing ALS. These include expansion mutations in the gene *C9orf72* (Dhasmana, Dhasmana et al. 2022) and exposure to low oxygen conditions (Vanacore, Cocco et al. 2010). However, genetic or environmental factors are rarely deterministic for developing ALS, but may, in combination, increase the risk. (Dharmadasa, Scaber et al. 2022). While the most prominent hallmark of ALS pathology involves the retraction of the axon from the neuromuscular junction, there is growing evidence of wider disruptions of mesoscale dynamics, i.e., changes in neuronal activity at the network and circuit level. The exact nature of these changes remains unclear, however signs of pathology appear early in the primary motor cortex and spread through interconnected networks to encompass broader cortical and subcortical brain structures as the disease progresses (Braak, Brettschneider et al. 2013). Furthermore, increased functional connectivity in affected brain networks has been reported for ALS patients (Douaud, Filippini et al. 2011) as well as asymptomatic carriers of predisposing genetic mutations (Menke, Proudfoot et al. 2016, Shoukry, Waugh et al. 2020) suggesting that brain network connectivity changes precede clinical symptoms. Furthermore, both these groups, as well as animal models of ALS, exhibit hyperexcitability and hyperconnectivity (Gunes, Kan et al. 2022). While ALS and underlying susceptibility to ALS have been shown to affect functional dynamics within networks of selectively vulnerable motor neurons, the mechanisms of these changes, including why such ongoing pathological changes do not overtly manifest as symptoms in the prodromal stages of the disease, remains unknown. A key challenge is the spatiotemporal aspect of evolving pathology within vulnerable motor neuron networks at the microscale and mesoscale, which complicates or entirely precludes studies in animal models or human patients.

However, we and others have shown that neural network dynamics at the microscale and mesoscale can be studied with advanced cellular models utilizing the inherent self-organizing properties of neurons into complex, computationally competent networks, akin to networks in the brain (Pasquale, Massobrio et al. 2008, Downes, Hammond et al. 2012, Poli, Pastore et al. 2015, Schroeter, Charlesworth et al. 2015, Fiskum, Sandvig et al. 2021, Antonello, Varley et al. 2022, Heiney, Huse Ramstad et al. 2022). Relevant applications of these methods to model ALS have been able to capture signs of hyperexcitability and time-dependent changes in networks of patient derived MNs (Wainger, Kiskinis et al. 2014, Ronchi, Buccino et al. 2021, Sommer, Rajkumar et al. 2022). However, investigations of more complex network dynamics are currently lacking. This pertains to both methodological and conceptual approaches applied to recapitulate, monitor, and decipher complex network dynamics. Approaches integrating connectomics and advanced neuroengineering are highly relevant in this context. Connectomics, with the application of graph theory, considers any network as a series of nodes connected by edges. In the brain or within a network of neurons, nodes can be single neurons, recording electrodes or voxels, and edges can be physical connections or temporal relationships. Using this approach, graph theory can derive descriptive measures from the complex interactions the networks produce (Bullmore and Sporns 2009). Application of graph theory to ALS patient magnetoencephalogram data has revealed trends of increasingly centralised networks which progressively rely on a few hyperconnected nodes to facilitate network function (Sorrentino, Rucco et al. 2018). Hyperconnectivity among a central group of network hub nodes, referred to as a rich club, has been proposed as a ubiquitous response to neuronal damage (Hillary and Grafman 2017). While supporting evidence is emerging for traumatic brain injury (Hillary, Rajtmajer et al. 2014), Alzheimer’s disease and multiple sclerosis (Hillary, Roman et al. 2015), the work by Sorrentino et al (Sorrentino, Rucco et al. 2018) is the only work showing increased brain network centralisation in ALS.

Self-organising *in vitro* neural networks mirror the behaviour of neurons and networks in the brain (Shein-Idelson, Ben-Jacob et al. 2011, Poli, Pastore et al. 2015, Schroeter, Charlesworth et al. 2015), and as such represent computational substrates. Connectomics and graph theory can thus be applied to establish the network’s “health” by inferring and assessing network features which facilitate high capacity for computation as well as deviations from such dynamics, as we have recently shown (Heiney, Huse Ramstad et al. 2021). By integrating these approaches with advanced neuroengineering and electrophysiology, we have also demonstrated that it is possible to probe micro- and mesoscale dynamic neural network reconfigurations as a result of (selectively) induced perturbations (Weir, Christiansen et al. 2023), including neurodegenerative pathology (Valderhaug, Ramstad et al. 2020, Valderhaug, Heiney et al. 2021).

In this study, we investigated the microscale and mesoscale dynamics of self-organising human ALS patient induced pluripotent stem cell (iPSC)-derived MNs harbouring an endogenous expansion mutation in *C9orf72*. We studied these networks over time both in high-density microelectrode arrays (MEAs), as well as custom designed multi-nodal microfluidic MEAs using complex network analysis of MEA electrophysiology data. MN networks derived from IPSCs with an expansion mutation in *C9orf72* organised into networks with hallmarks of high computational capacity to the same extent as healthy counterparts. However, they exhibited signs of dysfunction at the microscale and adaptations at the mesoscale consistent with existing hypotheses of hyperconnectivity and increased centrality as a response to neural network damage. This is the first study to describe functional connectivity changes in neuron networks derived from a patient with confirmed neurodegenerative disease. We further provide first-time evidence that endogenous connectivity and activity changes in ALS patient brain networks can be recapitulated in *in vitro* engineered neural networks, and that these properties cause the ALS networks to be more vulnerable to perturbation.

## Results

### ALS motor neurons exhibit endogenous TDP-43 proteinopathy, which is exacerbated after transient induced perturbation

Immunolabelling of MN networks derived from both an ALS patient with confirmed expansion mutation in *C9orf72* and healthy controls confirmed expression of MN markers Islet1, ChAT and HB9 (Supplementary figure 1). MNs also exhibited cytoplasmic TDP-43 inclusions, examples of which are shown in Figure 1A. Furthermore, immunolabelling of unperturbed motor neurons revealed that ALS patient derived MN networks exhibited more cytoplasmic inclusions of TDP-43 than healthy networks (p=0.0021, Wilcoxon rank-sum test) as seen in Figure 1B. Following induced transient hypoxia by means of cobalt chloride exposure, healthy networks, shown in Figure 1C, exhibited a transient rise in TDP-43 inclusions by 24 hours (p=0.0126, Wilcoxon rank-sum test), followed by a decline (p=0.0125, Wilcoxon rank-sum test) between 24 and 48 hours, back to baseline (i.e., pre-perturbation) levels. ALS patient derived networks, shown in Figure 1D, exhibited no significant changes in inclusions following hypoxic perturbation (all p>0.05, Wilcoxon rank-sum test). While the transient rise in TDP-43 inclusions in healthy networks resulted in no significant difference between the groups at 24 and 48 hours, the healthy networks again showed significantly lower levels of inclusions compared to ALS networks by 72 hours (p=0.0337, Wilcoxon rank-sum test), as seen in Figure 1B.

**Figure 1.**
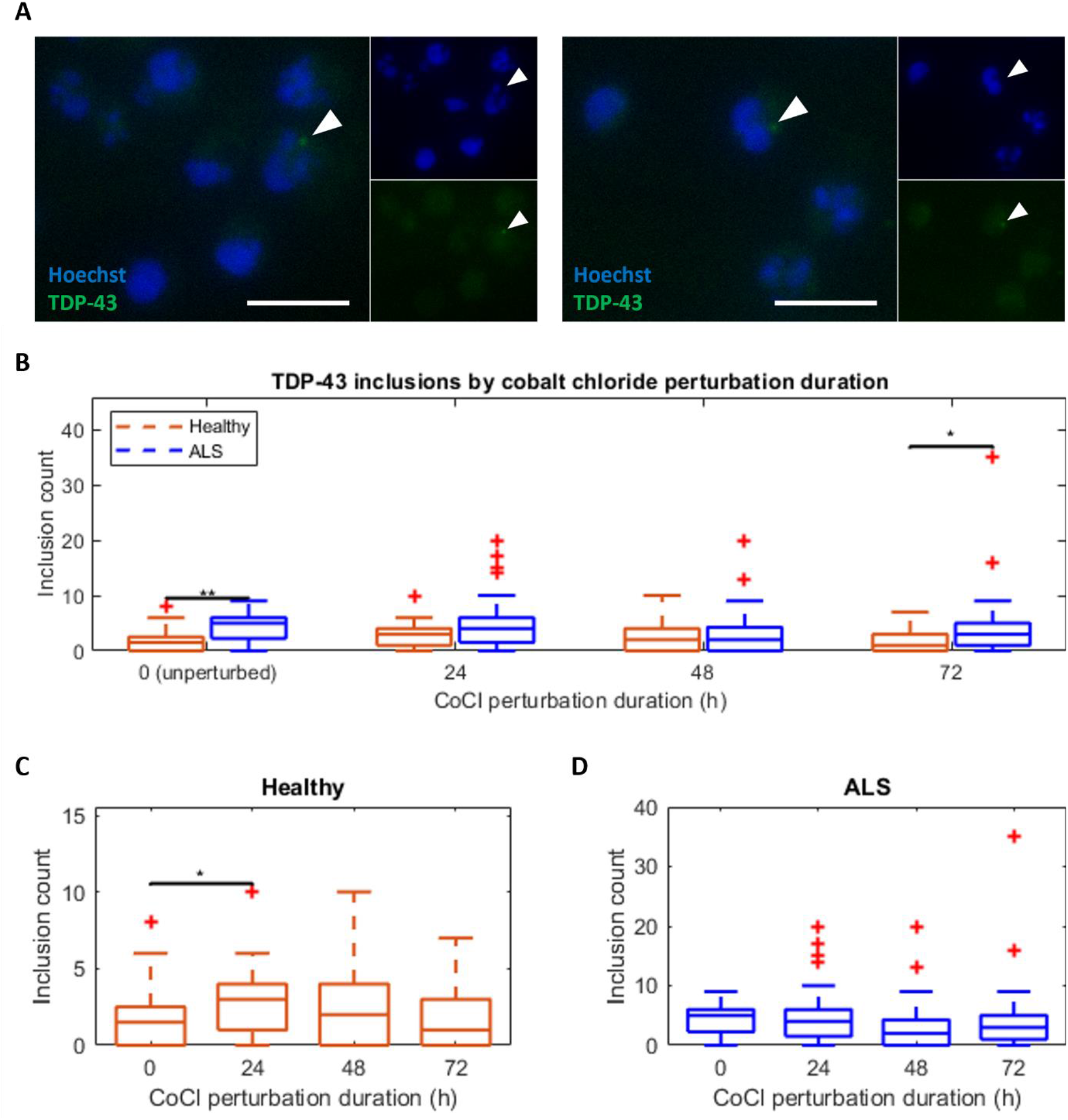
ALS motor neurons have endogenous TDP-43 proteinopathy, which is transiently exacerbated by cobalt chloride perturbation. A: White arrows highlight inclusions of TDP-43 (green) accumulating outside of the nucleus (blue) in ALS motor neurons. Scale bar = 25µm. B: ALS patient derived networks had a higher number of cytoplasmic inclusions of TDP-43 at 37 DIV, before CoCl perturbation, as well as 72 hours after perturbation. C: Healthy networks exhibited a transient rise in TDP-43 inclusion events following CoCl perturbation. D: ALS patient derived networks showed a small but not significant transient increase in TDP-43 inclusions following CoCl perturbation. Inclusions were counted in one microfluidic device per time point for both healthy and ALS conditions. Images examined per group: Healthy, perturbation duration=0h, n=28; perturbation duration=24h, n=27; perturbation duration=48h, n=30; perturbation duration=72h, n=26. ALS, perturbation duration=0h, n=23; perturbation duration=24h, n=28; perturbation duration=48h, n=29; perturbation duration=72h, n=27. *: p≤0.05, **: p<0.01, ***: p<0.001.

### ALS motor neurons self-organise into networks with hallmarks of high computational capacity

MN network activity from HD-MEAs was recorded every other day from 30 to 46 DIV. Figure 2A shows the HD-MEA, B an example of the voltage trace, C an example of the average spike waveform of a single electrode, D an example activity heatmap of a network and E an example connectivity map of the same network, where larger nodes have higher degree. Activity appeared stable throughout this period (see Supplementary figure 2). Group comparisons with GLMMs estimate a group average for healthy and ALS patient derived networks, which considers the dependent nature of the 9 repeated measurements of the same network. All results are reported by model estimated group average with 95% confidence interval (CI). Both healthy and ALS networks showed spontaneously emergent properties associated with highly computationally competent networks. This included similar levels of small-worldness, as seen in Figure 2G, where all networks had small world propensity > 0.6 (Healthy 0.822, CI [0.802-0.841]; ALS: 0.836, CI [0.816-0.855]). The node degree distributions of the networks adhered well to a power-law fitting, as seen in Figure 2H (Healthy 0.902, CI [0.894-0.910]; ALS: 0.911, CI [0.902-0.919]), which indicates that both healthy and ALS networks were scale free. Power law exponents were not significantly different between healthy and ALS networks (p=0.281, data not shown). Individual degree distributions are shown in Supplementary figure 4. The healthy and ALS networks also had similar degrees of modularity, shown in Figure 2I (Healthy: 0.292, CI [0.287-0.298]; ALS: 0.293, CI [0.288-0.299]). These traits are consistent with high computational capacity as outlined in (Heiney, Huse Ramstad et al. 2021). Lastly, although the estimated average network firing rate was higher in ALS networks, as seen in Figure 2F, this difference was not significant (Healthy: 834.4, CI [770.0-898.8]; ALS: 915.1, CI [850.7-979.5]).

**Figure 2.**
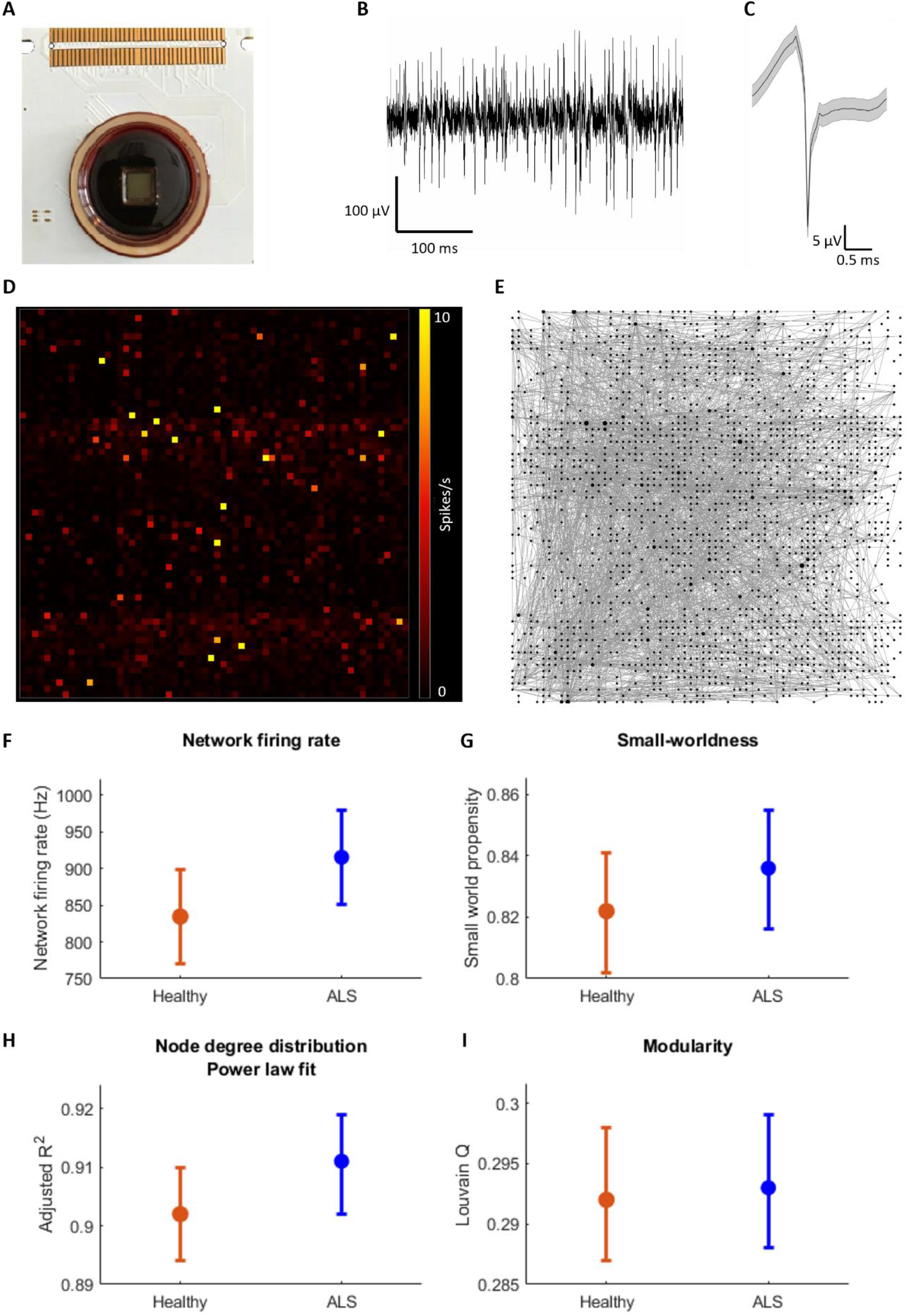
ALS patient derived and healthy motor neurons on HD-MEAs self-organise into networks with features of high computational capacity. A: High density MEAs allow for simultaneous sampling of neural activity from 4096 electrodes. B: Large deflections in the voltage trace recorded from the electrodes enables identification of likely action potentials. C: Following filtering, voltage deflections which exceed a certain threshold are identified as spikes. The waveform shows mean ± standard error of the spikes identified from one electrode. D: Identification of spikes from each electrode enables the construction of network activity maps. E: The temporal relationships between the firing of the electrodes enable the characterisation of functional connectivity and construction of graph maps. F-I show the results from generalised linear mixed models applied to 9 HD-MEA recordings per network (n=4 for both groups) in the period 30-46DIV. For daily results, see also *Supplementary* figure 2. F: Motor neuron networks derived from both a healthy donor and an ALS patient with an expansion mutation in the gene encoding *C9orf72* develop into networks with similar levels of activity. G: Healthy and ALS networks also develop similar degrees of small-worldness, H: Nodes in healthy and ALS networks both have similar degree distributions which follows a power law, indicating scale-free networks and I: Healthy and ALS networks both develop similar degrees of modularity. The similarity of the features shown in G-I indicate that healthy and ALS networks both self-organise into networks with high computational capacity. Plots show GLMM estimated group averages with 95% confidence intervals. No significant differences were found between healthy and ALS networks for these measures. *: p≤0.05, **: p<0.01, ***: p<0.001.

### ALS motor neuron networks exhibit microscale dysfunction and mesoscale compensation

Further analysis of the HD-MEA recordings from 30 to 46 DIV using GLMM revealed differences in multiple network features, as shown in Figure 3. Compared to healthy networks, ALS patient derived networks exhibited an average higher firing rate per electrode (Healthy: 0.090, CI [0.079-0.100]; ALS: 0.108, CI [0.098-0.119]; p=0.014), lower spike amplitudes (Healthy: 43.0, CI [42.4-43.7]; ALS: 41.5, CI [40.8-42.1]; p<0.001), as well as reduced electrode bursting (Healthy: 0.268, CI [0.250-0.286]; ALS: 0.219, CI [0.202-0.237]; p<0.001) compared to those of the healthy networks, all indicating microscale dysfunction in neuronal signalling. However, at the network level there was an opposite trend showing a higher fraction of spikes in network bursts (Healthy: 0.059, CI [0.036-0.082]; ALS: 0.092, CI [0.069-0.115]; p=0.049), while simultaneously exhibiting signs of less synchrony than healthy networks, as indicated by coherence index (Healthy: 0.083, CI [0.076-0.090]; ALS: 0.068, CI [0.061-0.075]; p=0.003). It is noteworthy that the ALS patient derived network had a significantly higher rich club coefficient compared to the healthy networks (Healthy: 0.611, CI [0.584-0.638]; ALS: 0.658, CI [0.631-0.685]; p=0.018). These features indicate mesoscale compensation involving increased dependency on a subset of highly interconnected nodes to maintain network functionality.

**Figure 3.**
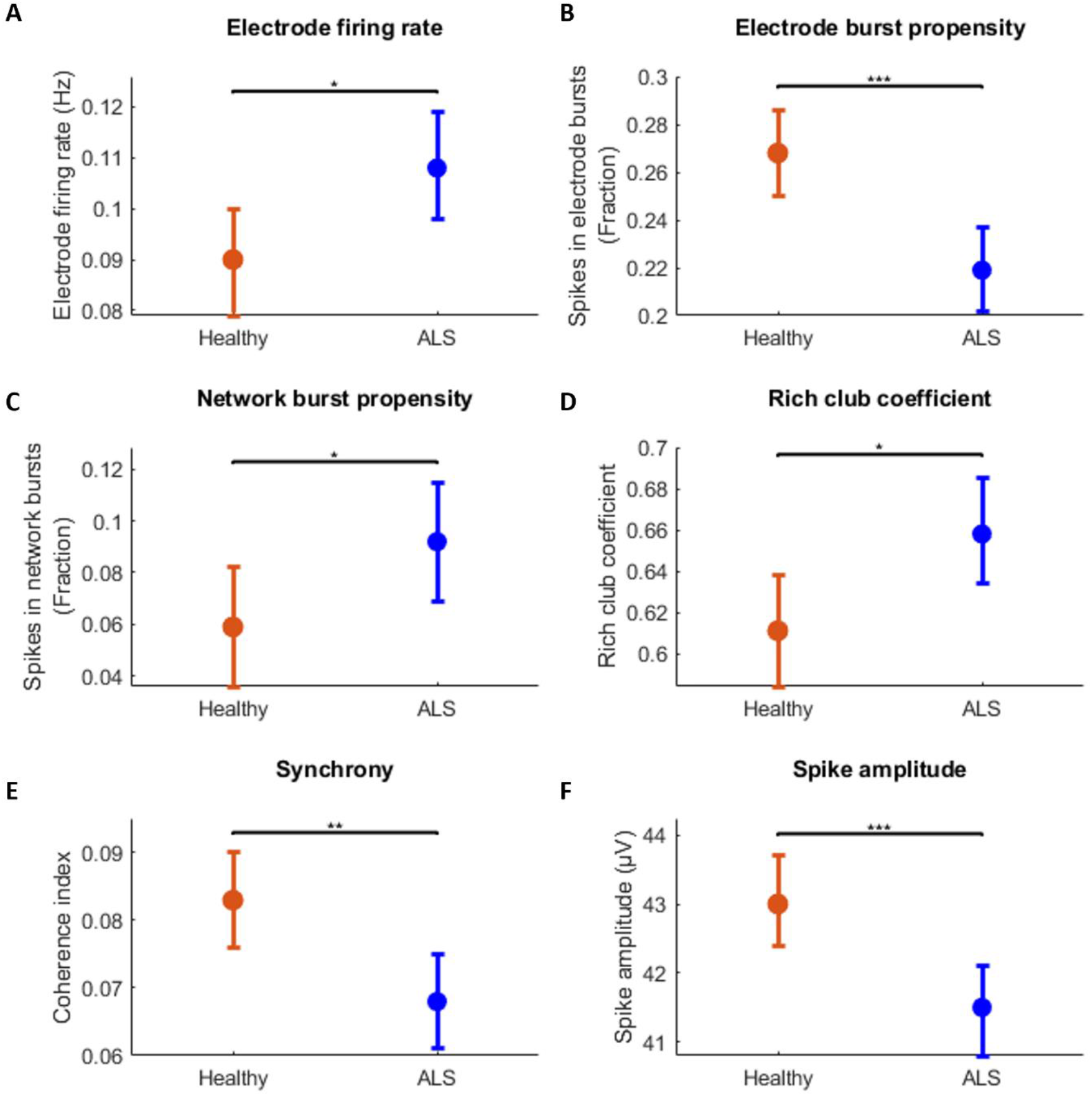
ALS motor neuron networks exhibit microscale dysfunction and mesoscale compensation consistent with increased metabolic cost. Results from generalised linear mixed models applied to 9 HD-MEA recordings per network (n=4 for both groups) in the period 30-46DIV. For daily results, *Supplementary* figure 3. Motor neuron networks derived from an ALS patient with an expansion mutation in *C9orf72* exhibit significant differences in many important network features, including A: hyperactivity at the electrode level and B: reduced electrode burst propensity, while C: network burst propensity is elevated. D: ALS networks have more developed rich club dynamics. E: Activity in ALS networks tend to be less synchronised than in healthy counterparts. F: spikes recorded from ALS networks have reduced amplitude compared to healthy networks. Plots show GLMM estimated group averages with 95% confidence intervals. *: p≤0.05, **: p<0.01, ***: p<0.001.

### ALS motor neuron networks are more susceptible to transient hypoxic perturbation

Prior to induced transient hypoxia by means of selective CoCl perturbation in the microfluidic chamber highlighted in magenta in Figure 4A, networks of healthy and ALS patient derived MNs showed similar levels of activity, as seen in Figure 4B (Healthy w/o CoCl: 70.4, CI [-5.05-145.9]; Healthy w. CoCl: 58.5, CI [-17.0-134]; ALS w/o CoCl: 134.7, CI [59.4-210]; ALS w. CoCl: 72.2, CI [-3.35-147.7]; all p>0.05). Following hypoxic perturbation, both perturbed and unperturbed ALS networks showed higher firing rates than the healthy networks, but only the perturbed ALS networks demonstrated a significant difference, as seen in Figure 4C (Healthy w/o CoCl: 32.4, CI [13.4-51.4]; Healthy w. CoCl: 34, CI [15-53]; ALS w/o CoCl: 64.8, CI [41.5-88.1]; ALS w. CoCl: 74.3, CI [55.3-93.3]; p=0.018 between ALS perturbed and healthy perturbed, p=0.015 between ALS perturbed and healthy unperturbed. p values are both corrected for multiple comparisons). Healthy networks, both unperturbed and perturbed, as well as unperturbed ALS networks, showed no significant differences (all p>0.05).

**Figure 4.**
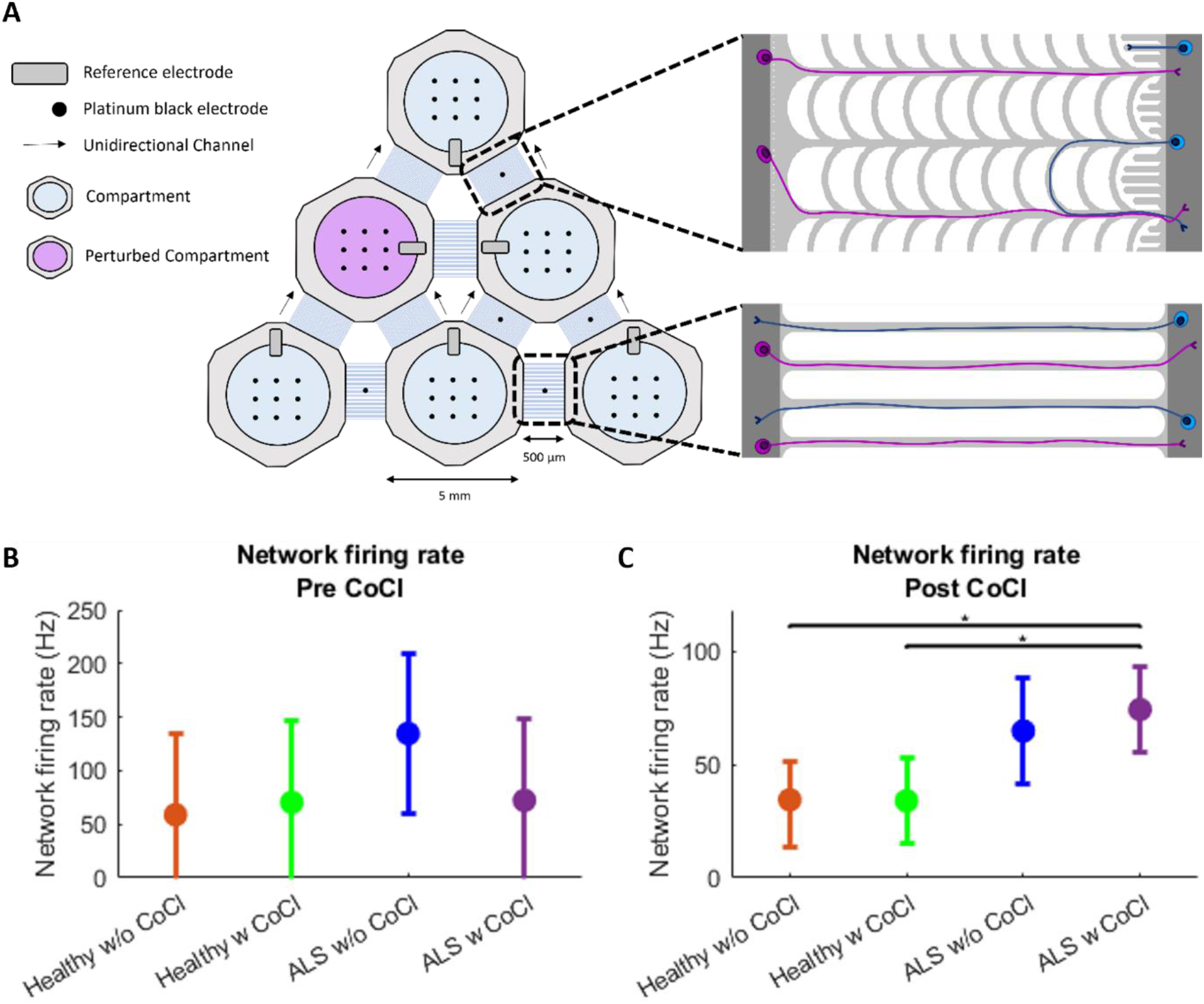
ALS motor neuron networks are more susceptible to transient hypoxic perturbation in microfluidic MEAs. A: The microfluidic multielectrode arrays were organised into 6 interconnected compartments with bidirectional (horizontal, bottom right) and unidirectional (indicated with arrows showing the directional selectivity, top right) channels. The compartment highlighted in magenta was selectively perturbed with cobalt chloride (CoCl). Microfluidic devices used for immunocytochemistry utilised the same design without the electrode interface. (b)-(c): Results from generalised linear mixed models applied to microfluidic multielectrode arrays in the period 30-47DIV. For daily results, see *Supplementary* figure 5. B: Prior to hypoxic perturbation, ALS and healthy networks showed no significant differences in overall network firing rate. C: Hypoxic perturbation caused hyperactivity in ALS networks relative to both unperturbed and perturbed controls. n=2 for ALS w/o CoCl, n=3 for all other groups. Plots show GLMM estimated group averages with 95% confidence intervals. *: p≤0.05, **: p<0.01, ***: p<0.001.

## Discussion

In this study we demonstrate for the first time that IPSC derived neural networks from a patient with confirmed neurodegenerative disease show endogenous pathological changes in functional connectivity. We show that ALS MN networks harbouring an expansion mutation in the gene *C9orf72* self-organise into networks with properties of high computational capacity to the same extent as healthy counterparts, as seen in Figure 2. However, we observed different underlying network dynamics, shown in Figure 3, that reveal increased centralisation and hub reliance in the patient-derived networks. These dynamics are consistent with higher overall network metabolic cost which makes the patient networks less resilient to perturbation despite the fact that they appear to be functioning normally. We identified clear signs of microscale dysfunction in ALS networks, including proteinopathy and functional activity consisting of weaker neuronal signalling, all consistent with increased metabolic cost and elevated network vulnerability. Specifically, we found that ALS patient derived MN networks had higher levels of TDP-43 cytoplasmic inclusions compared to controls, as seen in Figure 1. Cytoplasmic inclusions of TDP-43 are one of the most ubiquitous signs of MN cytopathology in ALS and frontotemporal dementia (Arai, Hasegawa et al. 2006, Neumann, Sampathu et al. 2006), and tend to be aggravated by increased oxidative stress, reduced antioxidant capacity, and mitochondrial dysfunction (Kara, Gordon et al. 2023). There is a substantial body of evidence supporting that MNs affected by ALS operate in a state of elevated oxidative stress (Cunha-Oliveira, Montezinho et al. 2020), and that environmental factors which further increase oxidative stress levels can contribute to risk and aggravation of the pathology (D’Amico, Factor-Litvak et al. 2013). We found that perturbation with increased oxidative stress through transient hypoxia induced by CoCl led to a temporary increase in cytoplasmic TDP-43 inclusions in healthy but not in ALS MN networks. 72 hours following perturbation and 24 hours after CoCl washout, the healthy networks had returned to a significantly lower level of inclusions than the ALS networks. Given that the levels of TDP-43 inclusions were higher both before perturbation and after perturbation, our findings further indicate that ALS networks operate at a higher baseline level of oxidative stress and metabolic cost.

MN hyperexcitability is a widely reported feature of ALS, but there are still unanswered questions with respect to network excitability dynamics during disease progression as well as their association with pathology. We found that ALS networks demonstrated microscale hyperactivity, with significantly increased firing rate at the electrode level, as seen in Figure 3A. Despite having higher levels of activity, we found that ALS networks had significantly less spikes occurring within bursts at the electrode level compared to healthy networks as seen in Figure 3B. Bursting consists of short periods of high frequency firing, which is important for robust information transfer between neurons (Zeldenrust, Wadman et al. 2018). Furthermore, it has been shown that reduced Ca^2+^ influx can reduce burst firing (Williams and Stuart 1999). It is therefore notable that spikes recorded from ALS networks had significantly lower voltage amplitudes than their healthy counterparts, as seen in Figure 3F. This is consistent with previous observations in ALS patient derived MN networks (Sommer, Rajkumar et al. 2022). Lower spike amplitude results in reduced Ca^2+^ influx (Scarnati, Clarke et al. 2020), and may represent an adaptation of ALS MNs to minimise Ca^2+^-induced excitotoxicity and oxidative stress. However, such functional dynamics also reduce the amount of neurotransmitter released as a consequence of the spike (Scarnati, Clarke et al. 2020), and may therefore contribute to disruption of the network’s function. We observed this in terms of the electrode burst propensity but also the overall synchrony of the network, which we found to be lower than in healthy networks, as seen in Figure 3E. Overall, at the microscale, we found that ALS MNs generate more, but weaker spikes, which leads to uncoordinated functional activity and less robust signalling, consistent with higher metabolic cost and increased oxidative stress.

Interestingly, despite these differences, we observed that similar to healthy MNs, ALS MNs self-organised into networks with hallmarks of high computational capacity, as shown in Figure 2. At the mesoscale level, both groups showed strong evidence of small world and scale free network dynamics, as well as similar levels of modularity. However, we found that despite having reduced burst propensity at the electrode level, ALS networks contained a higher proportion of spikes within network wide bursts compared to healthy networks, as seen in Figure 3C, indicating a more centralised signalling behaviour. Furthermore, when we investigated the functional connectivity in greater detail, we found that the ALS networks had a higher interconnectivity of high degree nodes, as evidenced by the elevated rich club coefficient in Figure 3D, indicating a more centrally connected network organisation. A rich-club is an interconnected community of high degree nodes within a network which facilitate global integration of signals across different parts of the network (Griffa and Van den Heuvel 2018). Increased connectivity within a rich-club appears to be a common response in neural networks exposed to damage (Hillary, Rajtmajer et al. 2014). When local connections are impaired, either by loss of nodes or loss of signal fidelity as seen in the reduced spike amplitude and electrode bursting in our findings, networks adapt by reorganising their functional connectivity. Because high degree hub nodes have high connectivity and activity, they are probabilistically more available for these adaptations, which therefore leads to an increased amount of information routing through hubs (Roy, Bernier et al. 2017). In an early phase of this adaptation, or if the network damage is limited to an acute perturbation that requires finite reorganisation, this may be beneficial for maintaining network communication (Hillary and Grafman 2017). However, such reorganisation increases the metabolic demand placed on hub nodes, and ongoing network damage such as in neurodegenerative disease leads to a continuous mounting stress on these nodes, which already represent some of the most metabolically costly components of the network (Bullmore and Sporns 2012)van den Heuvel, 2012 #422}. Eventually, the nodes succumb to this burden, and since so much of the network’s overall communication is dependent on them, the network fails to maintain its function without them (Stam 2014). While supporting evidence for such a pattern of damage and compensation has been found in cases of brain injury and Alzheimer’s disease (Hillary, Roman et al. 2015), it has not been strongly substantiated for ALS pathology until now. While the work by Sorrentino et al (Sorrentino, Rucco et al. 2018) shows increased centralisation in ALS patient brain networks, we expand on this by showing that increased connectivity is prominent within the rich club and that these networks are rendered highly vulnerable to perturbation. Our study thus provides clear evidence of pathological dynamics of centralisation occurring in ALS MN networks.

Our results from the microfluidic MEA-based assays showed that perturbation with CoCl had little effect on healthy network activity, but caused a rise in activity in ALS networks, as seen in Figure 4B-C. We observed that unperturbed ALS networks showed a trend of higher network activity compared to healthy networks, while the difference only reached statistical significance in the perturbed ALS networks. This indicates that differences in network dynamics are enhanced by externally induced perturbations, suggesting that such perturbations can exacerbate ongoing disease processes. Hypoxic perturbation as induced in this study has been shown to increase oxidative stress by the same pathways as environmental hypoxia (Cervellati, Cervellati et al. 2014). Moreover, oxidative stress and formation of reactive oxygen species are known contributors to ALS pathogenesis (Cunha-Oliveira, Montezinho et al. 2020). Further work using high-throughput tools has the potential to utilise such a vulnerability to assess progression of disease processes and the efficacy of interventions (Fiskum, Sandvig et al. 2021).

In this study we studied ALS MNs with an endogenous expansion in *C9orf72*, i.e., distinct from CRISPR-induced genetic predisposition or induced pathology by means of misfolded protein seeds. However, there are multiple other genes which increase the risk of developing ALS. Future studies should examine motor neuron networks derived from ALS patients with various genetic predispositions, as well as networks derived from patients with sporadic ALS, as well as asymptomatic donors who possess ALS-associated mutations, to assess if the network features observed are general trends of ALS or specific to certain patient groups. It is also possible that patients with the same identified genetic predisposition may not share the same network features due to other, factors including epigenetic imprints. In this case, IPSC based network models in combination with complex network analysis, and also with network models developed by cell reprogramming that preserves epigenetic signatures by bypassing the stem cell stage, can contribute to personalised medical assessment based on the findings in patient-specific MNs (Okano and Morimoto 2022). Lastly, the methods and principles utilised in this paper can be applied to model and assess network dysfunction in other neurodegenerative diseases and to uncover which features of neural network dynamics, if any, may be shared by multiple pathologies.

In conclusion, in this study we show, for the first time, that motor neuron networks from a patient with confirmed neurodegenerative disease exhibit endogenous functional network changes which reflect changes observed in multiple pathological conditions *in vivo* in addition to classical proteinopathy. We provide clear evidence of microscale dysfunction and mesoscale compensation in ALS MN networks, including new evidence of increased network centralisation. ALS networks also exhibit higher levels of cytoplasmic inclusions of TDP-43, showing that common pathological hallmarks of ALS are present within such networks even at early stages. At the microscale, i.e., at single electrode level, ALS networks exhibited hyperactivity despite weaker, less robust signalling. At the mesoscale level, the networks had increased network-wide recruitment and showed an increased connectivity within a rich-club. The shift to a more centralised network structure in the face of ongoing network damage has been shown in other neurodegenerative diseases but has not previously been clearly demonstrated in ALS. We show that the network shift to more centralised functional organisation is an adaptation that maintains network features of high computational capacity, but which renders the networks more susceptible to perturbation, as observed by hyperactivity in ALS networks following induced transient hypoxia. Taken together, we show for the first time that ALS MN networks with an endogenous expansion mutation in *C9orf72* exhibit pathological hallmarks of ALS and self-organize into centralised, computational competent networks with a highly connected rich club and elevated vulnerability to perturbation.

## Acknowledgements

This work was supported by the Olav Thon Foundation, ALS Norge, Alf Harborg’s fund, NTNU Enabling Technologies and The Research Council of Norway for the support to the Norwegian Micro- and Nano-Fabrication Facility, NorFab, project number 295864.

## Author contributions

VF formulated the research questions and planned the experiments and methodology with advice from AS and IS. NWH designed and fabricated the microfluidic devices and microfluidic multielectrode arrays and wrote the methods section “Microfluidic Platform Design & Fabrication” and the first paragraph of the methods section “Data analysis”. NC analysed the high-density multielectrode array data and wrote most of the methods section “Data analysis”. NWH performed blinded imaging of TDP-43 inclusions, while IS performed blinded counting of TDP-43 inclusions. VF carried out all other experimental work and wrote the rest of the manuscript, with input from NWH, NC, AS and IS.

## Declaration of interests

The authors declare no competing interests.

## Supplementary material

**Supplementary figure 1.**
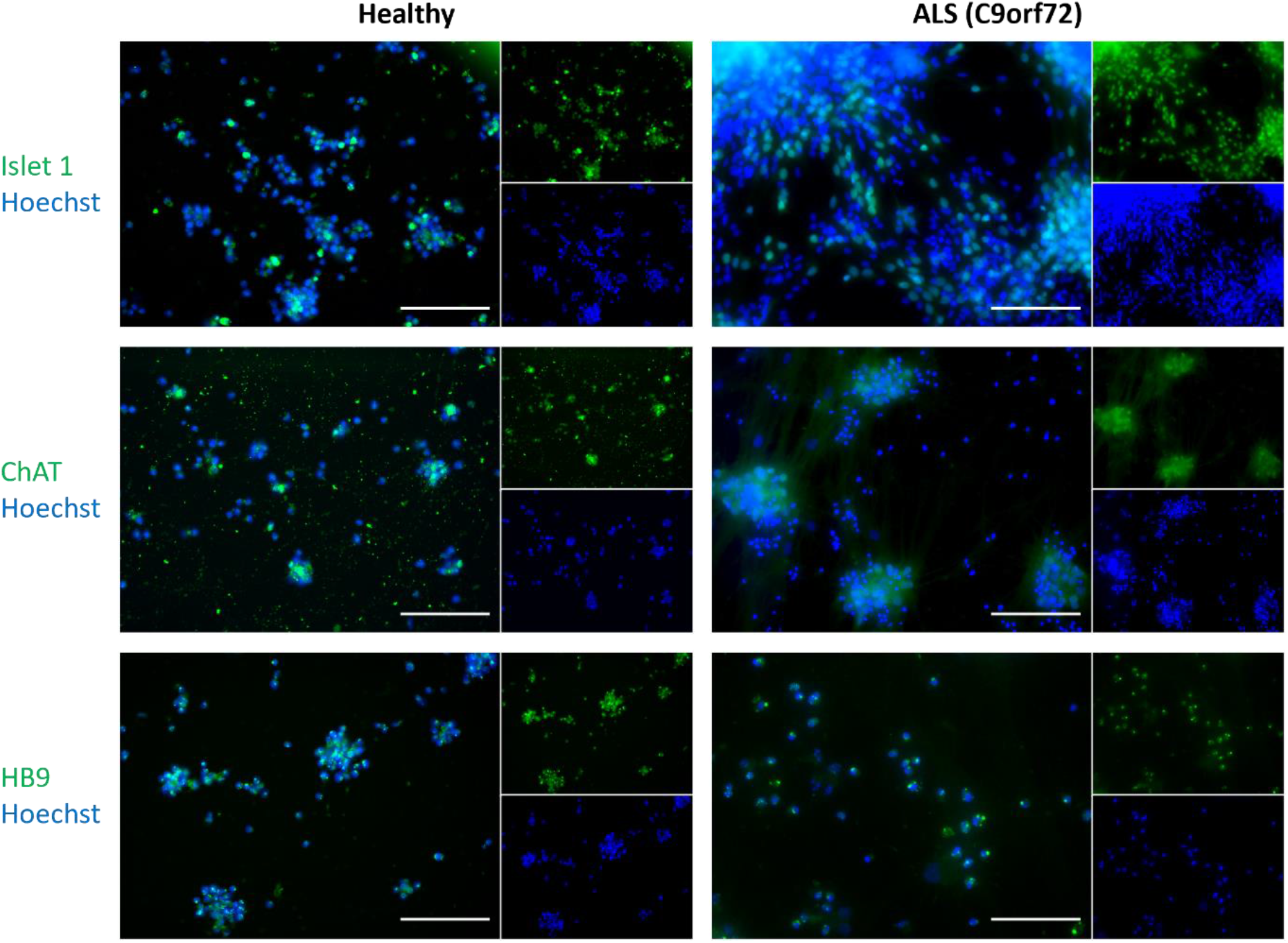
Immunocytochemistry confirms motor neuron cell identity. Both healthy and ALS (C9orf72) patient derived motor neuron networks showed expression of motor neuron specific markers Islet1, ChAT and HB9. Scale bar = 250 µm.

**Supplementary figure 2.**
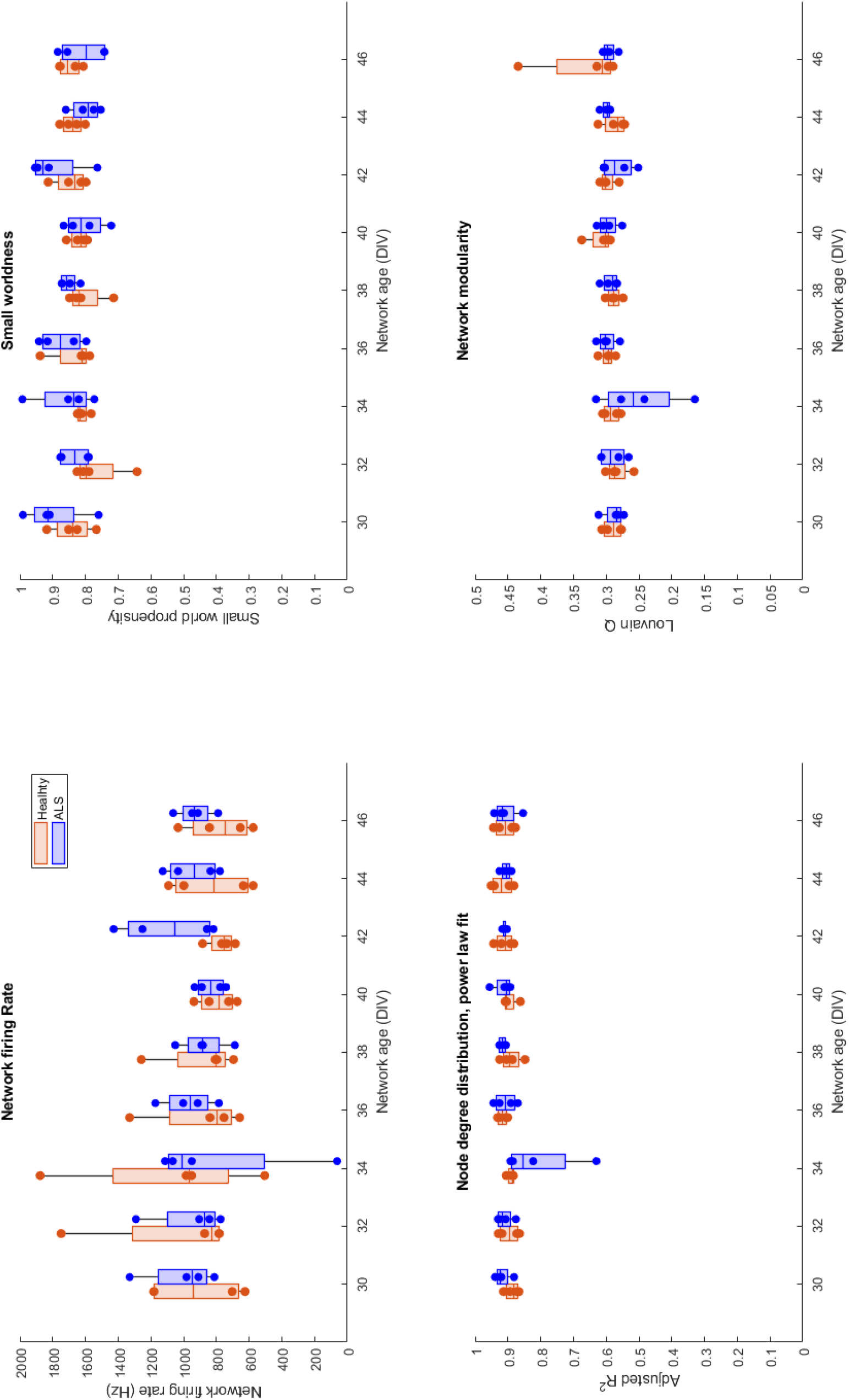
Development of motor neuron network activity on HD-MEAs; features of high computational capacity. Development of network activity and individual data points used to generate the GLMMs in Figure 2

**Supplementary figure 3.**
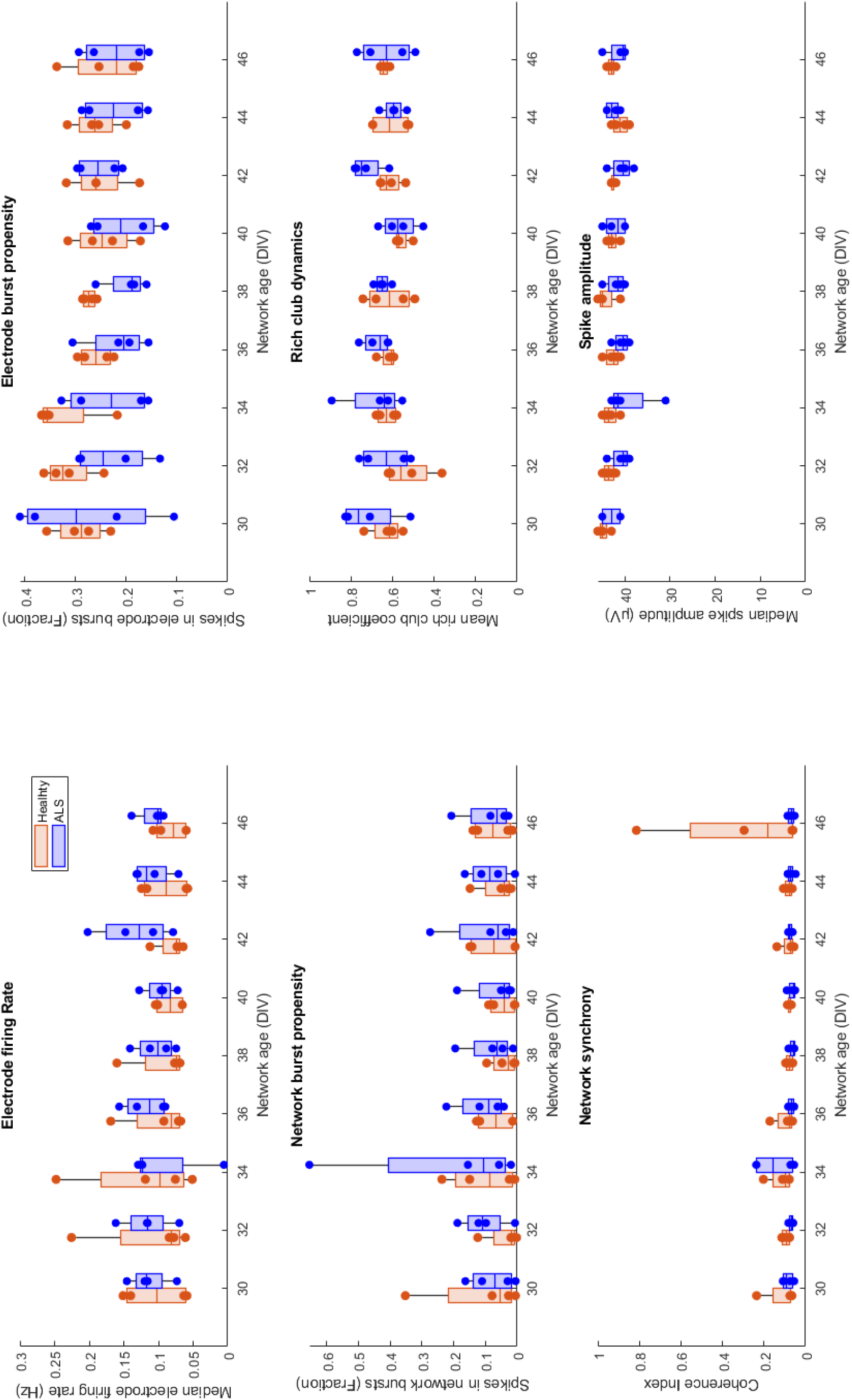
Development of motor neuron network activity on HD-MEAs; microscale dysfunction and mesoscale compensation. Development of network activity and individual data points used to generate the GLMMs in Figure 3.

**Supplementary figure 4.**
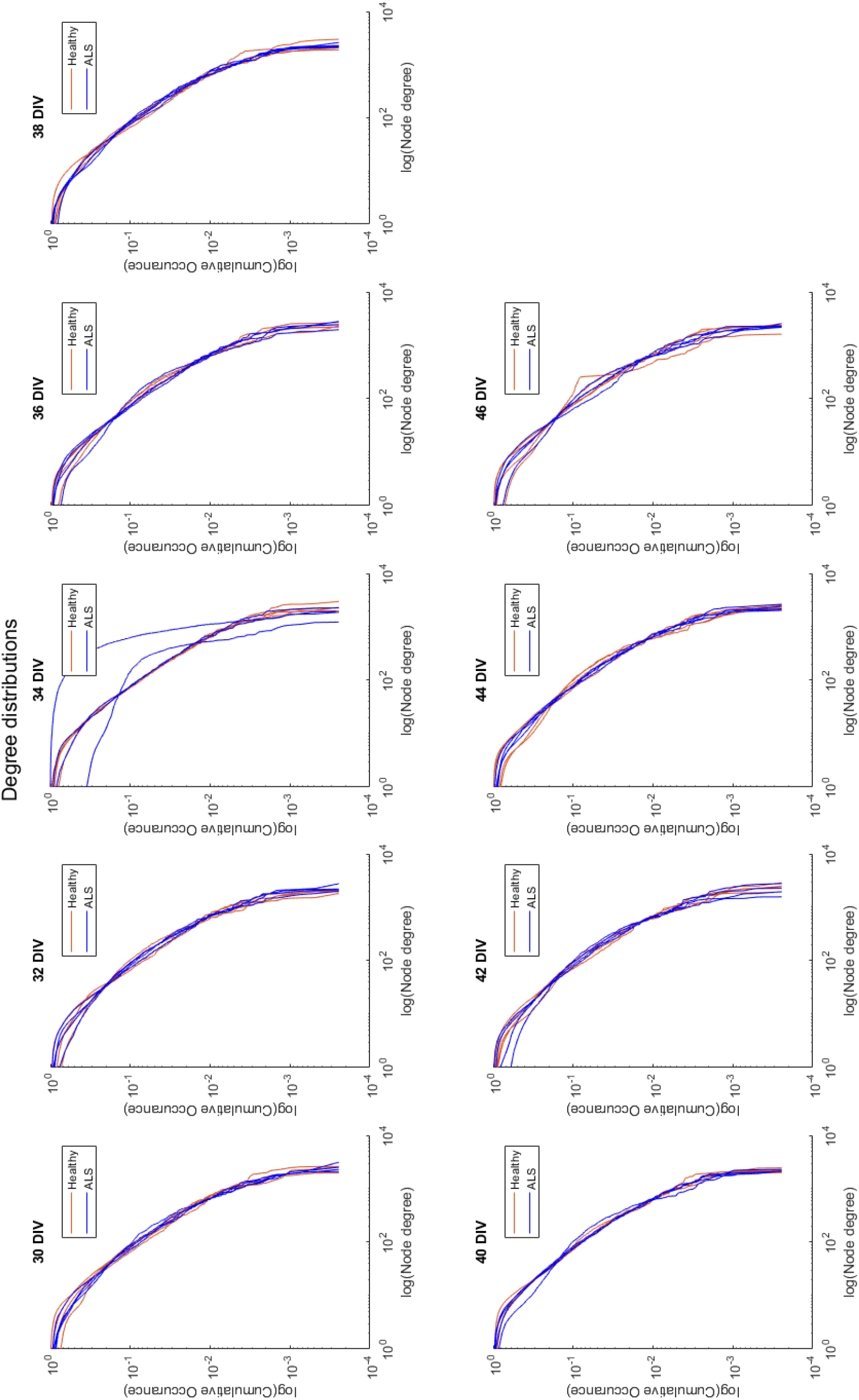
Motor neuron network degree distributions. Motor neuron networks on HD-MEAs have node degree distributions which appear to follow a power-law, i.e. linear in a log-log scale.

**Supplementary figure 5.**
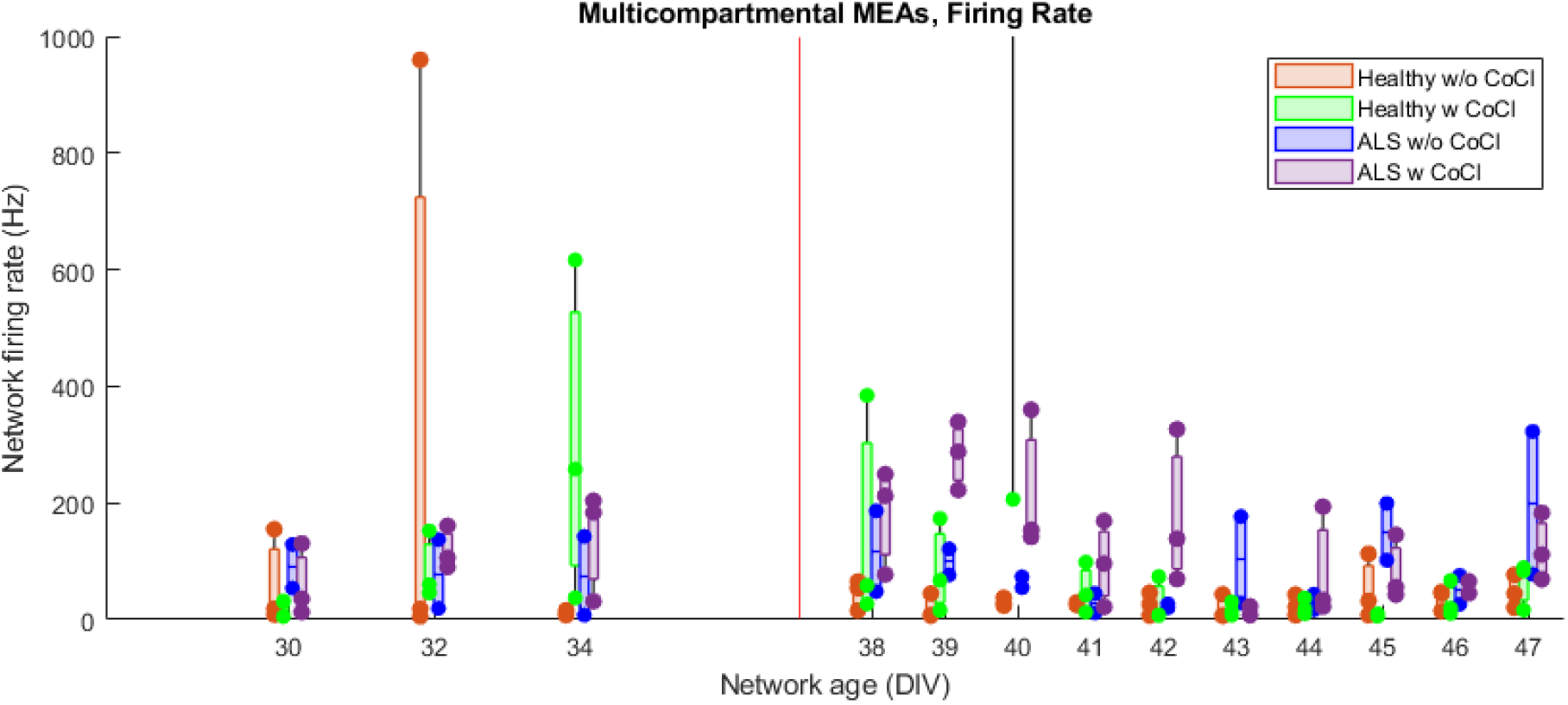
Development of network firing rate of motor neuron networks on microfluidic MEAs. Development of network firing rate and individual data points used to generate the GLMMs and longitudinal variance in Figure 4B-C. The red vertical line indicates DIV 37, when CoCl perturbation was applied. Two data points are not shown for 40DIV, Healthy control, Hypoxic due to large outlier values (3955.77 and 12375.4).

## Online methods

### Microfluidic Platform Design & Fabrication

Microfluidic platforms comprising six compartments were designed to structure MN networks with a hierarchical, modular structure. Furthermore, the design allowed for selective perturbation to parts of the network. Designs for the microdevices were created using the CAD software Clewin 4 (WieWeb Software, Enschede). A schematic of the design can be seen in Figure 4A. Each compartment was 5 mm wide and 60 mm high, employing a partially open design, initially demonstrated by van de Wijdeven *et al* (van de Wijdeven, Ramstad et al. 2018). Compartments within the same levels of the hierarchy were connected bidirectionally through 52 microchannels, each 500 μm long, 10 μm wide and 5 μm high. Compartments across the levels of the hierarchy were connected unidirectionally through 20 microchannels, each 700 μm long, 10 μm wide and 5 μm high. To promote unidirectional axonal outgrowth between compartments across the levels of the hierarchy, geometrical constraints inspired by the arched return-to-sender design of Renault *et al* (Renault, Durand et al. 2016) were incorporated in the microfluidic channels, effectively redirecting axons from the postsynaptic side to the chamber from which they originated. Furthermore, axon traps were incorporated on the postsynaptic side to prevent any outgrowing axons from reaching the presynaptic compartment. A pattern of 4 μm diameter pillars with 4 μm interspacing was positioned on the presynaptic side to prevent neuronal somata from entering and blocking the channels.

For immunocytochemistry (ICC), microfluidic devices (n=4 for both healthy and ALS conditions) were bonded to No. 1 glass coverslips (Menzel-Gläser, VWR International). For electrophysiological recordings, microfluidic devices were interfaced with microelectrode arrays (MEAs) (n=6 for both healthy and ALS conditions). 59 platinum electrodes of 100 μm diameter were positioned evenly spread across the array. 9 electrodes were centred in each compartment with 600 μm interspacing, while 5 electrodes were placed selectively in the channels to confirm active connections between the compartments. All electrodes had a thin layer of nanoporous platinum for enhanced signal-to-noise ratio. A reference electrode was furthermore split between the six compartments. A full protocol for the fabrication of the microelectrode arrays and microfluidic devices can be found in the preprint by Winter-Hjelm *et al*. (Winter-Hjelm, Tomren et al. 2022).

### Human iPSC culture and motor neuron differentiation

Human IPSCs were obtained from a healthy donor (female, 49 years) and a patient with confirmed ALS with *C9orf72* expansion (female, 64 years), both reprogrammed according to the Sendai virus IPSC reprogramming method. IPSCs were cultured and differentiated into MNs according to the protocol by Nijssen *et al*. (Nijssen, Aguila et al. 2019), specifically the section “Procedure: E” regarding MNs from human IPSCs, with the exception that an orbital shaker was used throughout the embryoid body phase. The timeline for MN differentiation and maturation is shown in Methods figure 1, and the composition of cell media at different stages is shown in Methods table 1.

**Methods figure 1.**
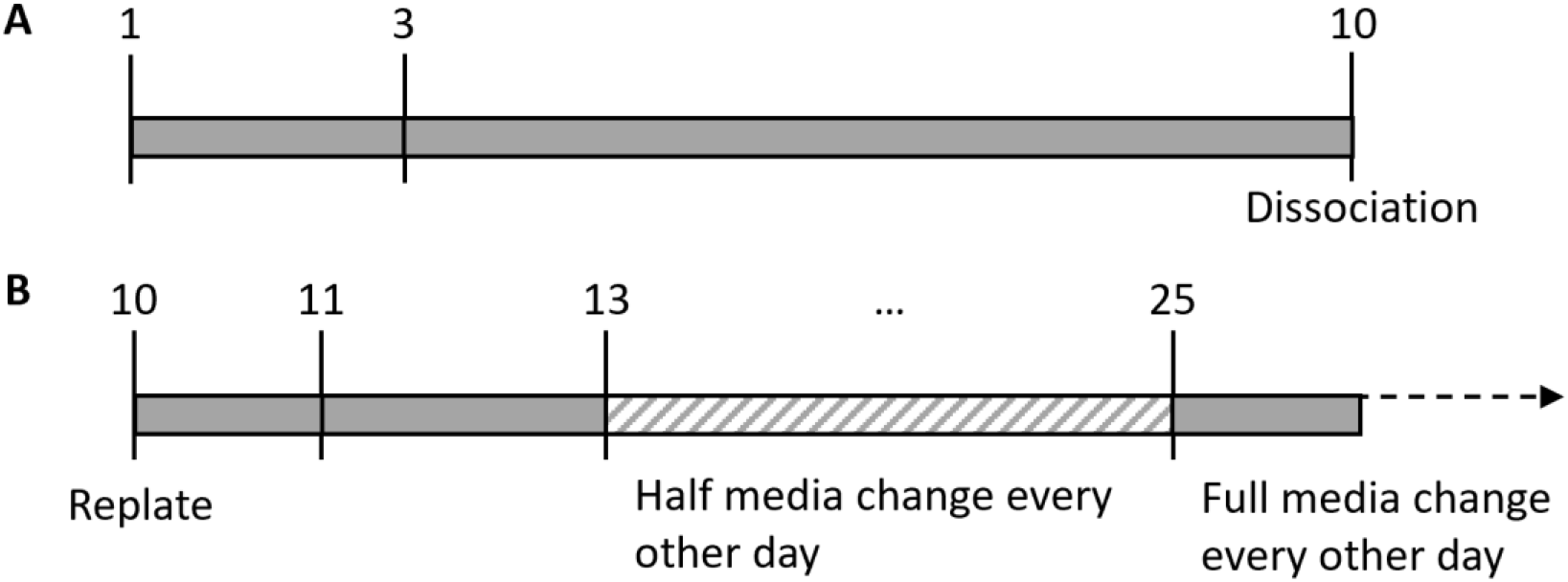
Timeline of motor neuron differentiation and maturation. A timeline showing the steps involved in the differentiation of motor neurons from induced pluripotent stem cells. The numbers indicate the day of the process, and vertical lines indicate a change in media composition or replacement regimen. A: During the initial phase, cells were maintained as embryoid bodies by using an orbital shaker. At the end of this period, embryoid bodies were dissociated prior to replating on the final vessels. B: After dissociation, single-cell suspensions were replated onto prepared vessels where the networks matured. The striped sections show a period when cell media was only half replaced every other day to minimise stress during maturation.

**Methods table 1.**
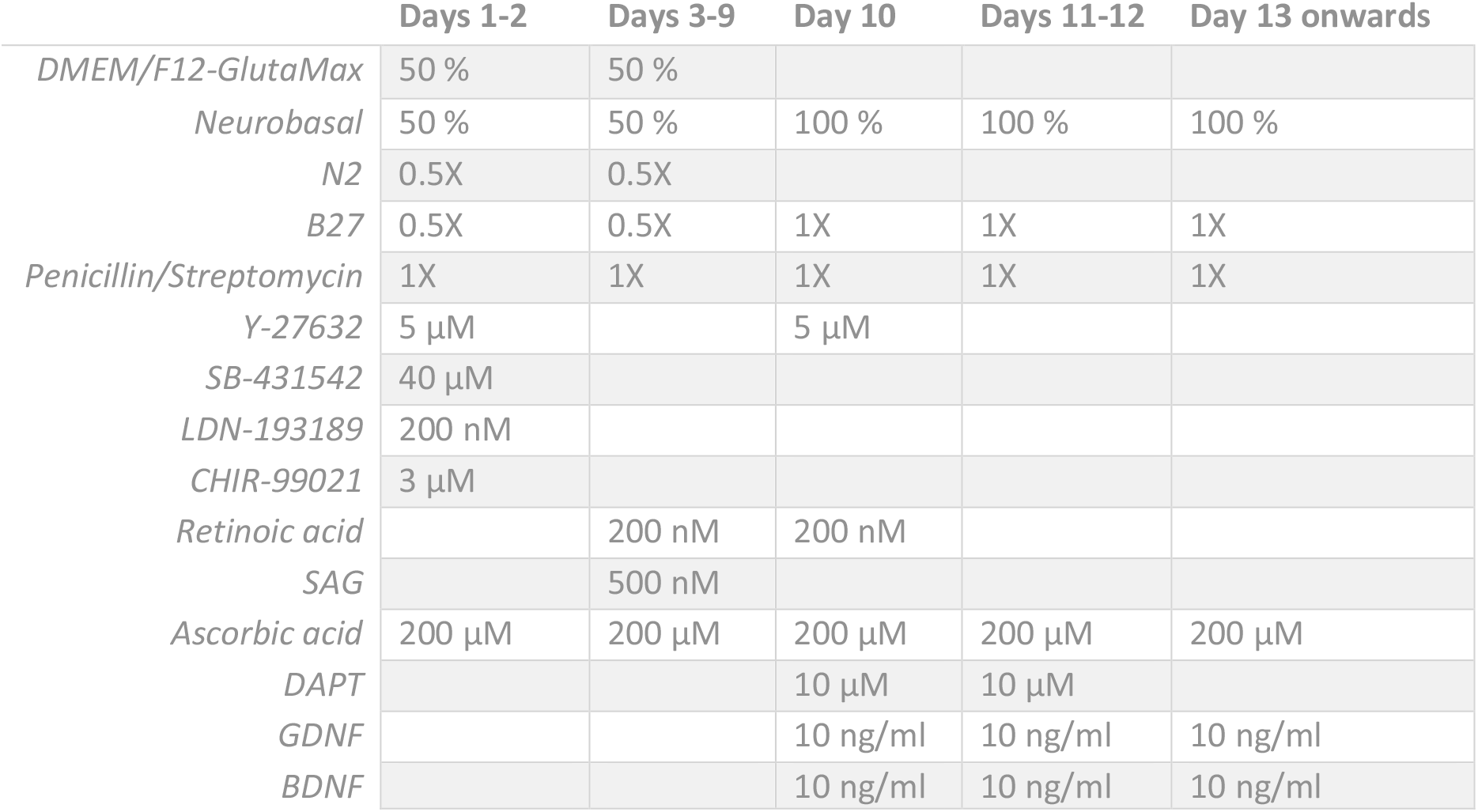
Cell media compositions during motor neuron differentiation. All cell media components were sourced according to Nijssen et al. (Nijssen, Aguila et al. 2019), and supplements were added to a base media composed of the indicated mix of DMEM/F12-GlutaMax and Neurobasal.

### Culture vessel coating, cell seeding and maintenance

Prior to seeding, the different culture vessels were coated to facilitate cell adhesion. HD-MEAs (3Brain Arena) with 4096 CMOS electrodes, n=4 for both healthy and ALS conditions, were first sterilised according to the manufacturer’s instructions and incubated with media overnight to increase surface hydrophilicity. Then, HD-MEAs and 8-well chambered slides (Ibidi 80807-90) were all coated with 0.05% poly(ethyleneimine) (P3141, Merck) diluted in HEPES (H0887, Merck) and incubated overnight. After rinsing four times with distilled water, they were left in room temperature to dry overnight in a laminar flow hood. Microfluidic devices were first sterilised for 12 hours under ultraviolet light, before coating with 0.01% poly-L-ornithine (P4957, Sigma) and incubated overnight. They were then rinsed four times with milli-q water for 15 minutes per cycle, ensuring different pressure gradients between chambers to rinse the channels. All vessels were then coated with 20 µg/mL natural mouse laminin (23017-015, Gibco) for 4 hours on HD-MEAs and 8-well chambered slides and 1 hour on microfluidic devices, before removing the laminin and seeding cells directly.

Cells were seeded at day 10 following the start of the MN differentiation protocol. Each HD-MEA was seeded with 80 000 live cells, each well in the 8-well chambered slides with 40 000 live cells and each well in the microfluidic devices and microfluidic MEAs with 15 000 live cells. After seeding, all vessels were incubated for 1 hour before filling with media, 1.5 mL in each HD-MEA, 400 µL in each 8-well chambered slide well and about 70 µL in each well in microfluidic devices and microfluidic MEAs. Media was fully (90%) replaced each day according to Nijssen *et al*. until day 13, shown in Methods figure 1B. After this point, media were half replaced every other day until day 25 to minimise stress to the networks during development. After this point, networks had become structurally stable, and medium was fully replaced every other day for the duration of the experiment.

### Electrophysiological recordings

For microfluidic MEAs, the average impedance of the electrodes was measured to 14.0 ± 8.3 kOhm in 0.1 % PBS solution using an MEA-IT60 System (Multichannel Systems). Network activity was recorded for 15 minutes using an MEA2100 system (Multichannel Systems) with a sampling rate of 20 kHz at 37°C. Prior to starting the recording, microfluidic MEAs were left to settle for 5 minutes. The activity of all networks was recorded every other day from day 30 to day 34, then every day from day 38 to day 47.

For HD-MEAs, network activity was recorded for 15 minutes using a Biocam Duplex System (3Brain) with a sampling rate of 18 857.72 Hz at 37°C. Prior to starting the recording, HD-MEAs were left to settle for 5 minutes. The activity of all networks was recorded every other day from day 30 to day 46.

### Cobalt chloride perturbation to induce chemical hypoxia

Cobalt chloride (CoCl) (15862, Merck) was used to induce chemical hypoxia (Zou, Yan et al. 2001) to selectively perturb one of the compartments in the microfluidic devices and microfluidic MEAs. At day 37, when networks had reached a stable state of activity, 1000 µM of CoCl were added to the middle-left compartment highlighted in magenta in Figure 4A. Half of all networks were perturbed, while the other half were left unperturbed (Perturbed healthy, n=3; unperturbed healthy, n=3; perturbed ALS, n=3; unperturbed ALS, n=2). When replacing media at 37 DIV, before CoCl was added, special care was taken to ensure the amount of cell media in the perturbed compartment was reduced relative to the other compartment in order to prevent diffusion of the CoCl to other compartments. The networks were maintained with this concentration of CoCl until the next media change 48 hours later at 39 DIV, which should be sufficient to induce widespread cell damage and death by chemical hypoxia (Zou, Yan et al. 2001). For motor neuron networks on microfluidic devices, three were perturbed with CoCl while one was left as an unperturbed control. After the perturbation, the three perturbed networks were individually fixed with glyoxal (see below) after 24, 48 and 72 hours respectively in order to assess the effect of the perturbation over time. The unperturbed network was fixed at 40 DIV.

Activity from MN networks on microfluidic MEAs was recorded every other day between 30 and 34 DIV. Transient hypoxic perturbation was induced at 37 DIV, after which network activity was recorded daily until 47 DIV (see Supplementary figure 5). Because of the hierarchical modular microfluidic design shown in Figure 4A, it was possible to introduce a selective perturbation to parts of the network and to assess the response of the entire network. Any residual CoCl was removed when cell media was replaced at 39 DIV. Overall network activity was considered before and after CoCl perturbation using GLMM.

### Immunocytochemistry

Immunocytochemistry (ICC) was applied to confirm MN identity after differentiation and maturation of the IPSCs and to investigate the presence of cytoplasmic inclusions of TDP-43. The approach was based on the work by Richter *et al*. (Richter, Revelo et al. 2018). Cell media was removed, and the networks were fixed in 3 % glyoxal solution for 15 minutes. Then, the networks were washed with PBS (D8662, Merck) four times. The cells were then permeabilised with 0.5 % Triton-X (T8787, Merck) in PBS for 5 minutes in 8-well chambered slides and with 0.2 % Triton-X for 10 minutes in microfluidic devices. Networks were then washed three times with PBS, before adding a blocking solution of 5 % goat serum (PCN5000, Fisher Scientific) in PBS for 1 hour at room temperature on an orbital shaker at 30 rpm. After aspirating the blocking solution, primary antibodies in PBS with 5 % goat serum were added and left overnight on a shaker table at 4°C. The next day, primary antibodies were removed, and networks were washed four times with PBS. Secondary antibodies were then added in PBS with 5 % goat serum and left on an orbital shaker at 30 rpm for 3 hours, followed by nuclear staining for 10 minutes. The networks were then washed four times with PBS, before being washed once with milli-q water. The duration of each washing step was 5 minutes in 8-well chambered slides and 15 minutes in microfluidic devices. Importantly, only 90 % of the PBS was removed in each washing step to reduce sheer stress on the network and prevent detachment. All 8-well chambered slides were fixed at 40 DIV, while microfluidic devices were fixed at different time points as described above.

MN networks were examined for expression of Islet-1 (ab109517, 1:250, Abcam), Chat (ab178850, 1:500, Abcam), HB9 (ab221884, 1:200, Abcam) and TDP-43 (PA5-27221, 1:500, Fisher Scientific), all visualized with the same secondary antibody (A11008, 1:1000, Fisher Scientific) and stained for Hoechst (62249, 1:2000, Fisher Scientific). Microfluidic devices were examined for cytoplasmic inclusions of TDP-43 by manual examination for labelled TDP-43 which did not overlap with Hoechst stain, as seen in the examples in Figure 1A. After fixation and immunolabelling, the identity of each microfluidic device was anonymized, and an independent investigator blinded to the study acquired 5 non-overlapping images per well for each microfluidic device. The identity of each image was then anonymized before a third independent investigator blinded to the study counted events of cytoplasmic inclusions of TDP-43 to allow comparison between healthy and ALS networks and the effect of CoCl perturbation. All images were acquired using an EVOS M5000 microscope with the following light cubes: DAPI (AMEP4650) and GFP (AMEP4651), and the following lenses: Olympus UPLSAP020x, 20x / 0.75 NA (N1480500) and Olympus UPLSAPO40x2, 40x/0.95NA (N2246700). All processing of the images was performed in Fiji/ImageJ.

### Data analysis

For microfluidic MEAs, raw data was filtered using a 4^th^ order Butterworth bandpass filter, removing high frequency noise (above 3000 Hz) and low frequency fluctuations (below 300 Hz). A notch filter was used to remove 50 Hz noise caused by the power supply mains. Both filters were run using zero-phase digital filtering using the Matlab 2020b (MathWorks) function filtfilt to avoid changes in the relative position of the detected spikes. Spike detection was conducted using the Precise Timing Spike Detection (PTSD) algorithm developed by Maccione *et al*. (Maccione, Gandolfo et al. 2009). The threshold was set to 8 times the standard deviation of the noise, the maximum peak duration to 1 ms and the refractory time to 1.6 ms.

For HD-MEAs, raw data was filtered using a 5^th^ order Butterworth highpass filter, removing low frequency noise (below 200Hz). Spike detection was conducted using the PTSD algorithm (Maccione, Gandolfo et al. 2009). The threshold was set to 8 times the standard deviation of the noise, the peak lifetime period duration to 1.5 ms and the refractory time to 1 ms. Filtering and spike detection was performed in BrainWave 5 (3Brain).

The electrode firing rate was determined by the number of spikes recorded for a single electrode divided by the total recording time, then taking the median of each network. The network firing rate was the total number of recorded spikes divided by the total recording time.

Bursts were defined as a sequence of at least four spikes with an inter-spike interval of less than 100 ms between each consecutive spike, performed for each electrode. Network bursts were defined as a sequence of at least 100 spikes within 12.5 ms, corresponding to the minimum between the bimodal peak of the inter-spike intervals between 100 spikes in the recording (Bakkum, Radivojevic et al. 2013). Evaluations of the minima were done manually and kept constant for all recordings. The fraction of spikes in bursts, or burst propensity, was calculated as the number of spikes in bursts divided by the total number of spikes and applied to both electrode and network bursts respectively. Coherence index was used as a relative measure of network synchrony (Wang 2002).

To generate the network connectivity matrix of the HD-MEA recordings, spike times were separated into 100 ms bins. Then, the co-occurrences of the binned spike times were identified, resulting in a count of the number of times spikes on different electrodes co-occur. Stronger connections will have a higher tendency to co-occur. To establish a threshold for non-spurious connectivity, randomized series of corresponding data were generated by shuffling the original data. This was repeated 10 times, and co-occurrence counts which were equal to or lower than the mean of the shuffled data were removed from the connectivity matrix. Of these connections, the 1 % of strongest connections were further selected, resulting in binary connectivity matrixes with a comparable number of connections. Finally, the giant component from this was analysed using common network metrics. Small world propensity provides an estimate to how well each network conforms to small world principles (I.e. high clustering and low average path length), as described in (Muldoon, Bridgeford et al. 2016). A small world propensity above 0.6 is considered small world. The rich club coefficient measures the mean rich club coefficient, which is a measure of the tendency for nodes with high degree to interconnect (McAuley, da Fontoura Costa et al. 2007). Community detection was done using CDlib (Rossetti, Milli et al. 2019), while modularity evaluation was done using NetworkX (Swart 2008) according to (Clauset, Newman et al. 2004). The Matlab function fit, with fitType=‘power1’ was used to assess if node degree distribution followed a power law (Matlab 2020b, MathWorks).

Generalised linear mixed models (GLMMs) were used to assess the differences in various network properties between healthy and ALS patient-derived MN networks. The models used genotype as a fixed effect with various network features as targets, utilising a linear model with repeated measurements of each network (the subjects of the model) and sequential Bonferroni adjustment for multiple comparisons. All GLMMs were implemented in SPSS version 29.0.0.0.

## Notes

### Competing Interest Statement

The authors have declared no competing interest.

